# Cardiomyocyte transcriptomic signatures in response to *Trypanosoma cruzi* infection underpin Chagas cardiomyopathy progression

**DOI:** 10.1101/2023.02.27.530371

**Authors:** Katherine-Sofia Candray-Medina, Yu Nakagama, Masamichi Ito, Shun Nakagama, Evariste Tshibangu-Kabamba, Norihiko Takeda, Yuki Sugiura, Yuko Nitahara, Yu Michimuko-Nagahara, Natsuko Kaku, Yoko Onizuka, Carmen-Elena Arias, Maricela Mejia, Karla Alas, Susana Peña, Yasuhiro Maejima, Issei Komuro, Junko Nakajima-Shimada, Yasutoshi Kido

## Abstract

Chagas disease can lead to life-threatening cardiac manifestations that occur more frequently in geographic areas more prevalent with the TcI/II circulating genetic strains. To elucidate the differential transcriptomic signatures of the cardiomyocyte resulting from infection with TcI/II or TcVI *T. cruzi* strains and explore their relationships with pathogenesis, HL-1 rodent cardiomyocytes were infected with TcI/II or TcVI *T. cruzi* trypomastigotes. RNA was isolated serially post-infection for microarray analysis. Enrichment analyses of differentially expressed genes (fold-change ≥2 or ≤ 0.5) highlighted the over-represented biological pathways. We found that Oxidative stress-related GO terms, ‘Hypertrophy model’, ‘Apoptosis’, and ‘MAPK signaling’ pathways (all with p<0.01) were upregulated. ‘Glutathione and one-carbon metabolism’ pathway, and ‘Cellular nitrogen compound metabolic process’ GO term (all with p <0.001) were upregulated exclusively in the cardiomyocytes infected with the TcI/II strains. Upregulation in the oxidative stress-related and hypertrophic responses are shared hallmarks with viral myocarditis, another inflammatory cardiac pathology. Nitrogen metabolism upregulation and Glutathione metabolism imbalance may implicate the relation of nitrosative stress and poor oxygen radicals scavenging in the unique pathophysiology of chagasic cardiomyopathy development.

**Importance:** Chagas disease affects more than 6 million people worldwide. One-third of those chronically infected will develop the life-threatening condition Chagas Cardiomyopathy (CCM). *Trypanosoma cruzi (T. cruzi)*, grouped based on their genetic variability into six discrete typing units (DTU), are associated with DTU-specific clinical phenotypes. The diverse genetic make-up of parasite virulence factors shall evoke unique host defense responses of variable magnitude, collectively affecting the phenotypic expression of CCM. To address this, we performed a transcriptome analysis of cardiomyocytes infected with three different *T. cruzi* strains each belonging to a different DTU. As a result, we were able to point out dysregulation in nitrogen metabolic processes, Glutathione, and one-carbon metabolism pathways as main features in the host response against cardiomyopathy-prone *T. cruzi* strains. Further research on these pathways could serve not only in the lookout for progression biomarkers but also in the lead toward the discovery of new therapeutic targets.

## Introduction

Chagas disease, caused by *Trypanosoma cruzi* (*T. cruzi)* protozoan infection, affects more than 6 million people worldwide with more than 70 million at risk of becoming infected (1–3). The clinical outcome of Chagas disease is highly variable, where only one-third of those with chronic infection develop overt Chagas cardiomyopathy (CCM), a life-threatening condition characterized by myocardial dilatation, conduction abnormalities, and heart failure (4) The molecular pathophysiology and indicators of CCM progression have been least described among other common forms of dilated cardiomyopathy, such as those related to ischemic/hypoxic stress (mitochondrial dysfunction and impaired calcium sensitivity of the cardiomyocyte) or viral infections (viral RNA persistence and subsequent apoptotic loss of the cardiomyocyte) (5–15).

The genetic diversity of *T. cruzi* has gathered attention as a determinant of Chagas disease phenotypic expression and CCM progression. *T. cruzi*, grouped based on their genetic variability into six discrete typing units (DTU), TcI to TcVI, are associated with DTU-specific clinical phenotypes (16,17). TcI -and less often, TcII-parasites have been most strongly associated with CCM (18–22), while TcVI-infected individuals more commonly present with the digestive-form disease (23,24). The diverse genetic make-up of parasite virulence factors shall evoke unique host defense responses of variable magnitude, collectively affecting the phenotypic expression of CCM. Thus, the host response against specific *T. cruzi* genotypes shall provide yet-to-be-clarified clues to the molecular pathophysiology of CCM progression.(25–27) The objective of the current study was to elucidate the differential transcriptomic signatures of the cardiomyocyte response against infection with variable *T. cruzi* strains, the CCM-prone TcI/II, or the less typical TcVI and explore the biological processes underpinning CCM progression.

## Materials and methods

### Cell lines and culture conditions

Mouse CH3/10T1/2, clone 8 murine fibroblasts (RRID: CVCL_0190, JCRB0003), obtained from the Japanese Collection of Research Bioresources Cell Bank (JCRB), were cultured in high glucose DMEM (Fujifilm, Japan) supplemented with 10% fetal bovine serum and 1% penicillin/streptomycin (Fujifilm, Osaka, Japan). Cells were seeded on T-25 vent-cap flasks and incubated at 37 °C, under 5% CO_2_.

Mouse HL-1 cardiomyocytes (RRID: CVCL_0303) were cultured in Claycomb medium (Merck, Darmstadt, Germany) supplemented with 10% fetal bovine serum (Gibco, Maryland, USA), 0.1 mM Norepinephrine (Sigma, Missouri, USA), 0.2 mM L-Glutamine (Fujifilm, Osaka, Japan), and 1% penicillin/streptomycin (Fujifilm, Osaka, Japan). Cells were seeded on T-25 vent-cap flasks pre-coated with fibronectin (Sigma, Missouri, USA) and incubated at 37 °C, under 5% CO_2_.

*T. cruzi* epimastigotes of the Colombian, Y, and Tulahuen strains (from TcI, TcII, and TcVI DTUs, respectively) were obtained from NEKKEN Bio-Resource center (NBRC) and cultured in LIT medium at 28 °C, under 5% CO_2_. Per strain, 1×10^8^ epimastigotes harvested in the exponential growth phase were subjected to metacyclogenesis following a pre-established protocol. (28) Debris sedimentation was performed with a centrifugation step at 700 g for 5 min followed by trypomastigote collection of the supernatant at 1700 g for 10 minutes. Trypomastigotes were used to infect CH3/10T1/2 cells with a multiplicity of infection (MOI) 1:10. After 5-7 days, the successful infection could be confirmed by the presence of intracellular amastigotes on light microscopy. Cultured-cell-derived-trypomastigotes (CCdT), obtained from the supernatant of these stable cultures through the aforementioned centrifugation condition, were used for CCM modeling experiments.

### *In vitro* CCM modeling

6-well plates were seeded with HL-1 cardiomyocytes as described before and incubated for 24 hours to ensure full attachment and confluency. CCdT of each strain were collected to infect HL-1 cardiomyocytes at a MOI of 10:1. 100 ng/mL of mouse TNFα (Sigma-Aldrich H8916, Missouri,USA) was added to the culture medium of positive control wells. Negative control cells underwent medium replacement. Each condition was prepared in duplicates.

### Microarray analysis of differentially expressed genes (DEGs)

At 24- and 48-hours post-infection (hpi), culture medium was removed, and cells were gently washed with sterile PBS. Total RNA was isolated with the NucleoSpin RNA kit (Macherey-Nagel, Düren, Germany) as instructed by the manufacturer. RNA concentration and purity was assessed with Nanodrop 2000 (ThermoFisher, Wilmington, USA). RNA samples were analyzed on the Clariom S Assay for mouse (Applied BioSystems, CA, USA) microarray platform to identify DEGs.

Data analysis was done on the Microarray Data Analysis Tool software version 3.2.0.0 (Filgen, Nagoya, Japan). Genes with signals below the threshold of 43, 48, 43, 52 at 24 and 48 hours on the T cruzi-infected (CCM model) and control conditions, respectively, were excluded leaving 10,784 valid gene probes per condition for analysis. Fold change (FC) was defined as the ratio between gene expression level of the CCM models (or the positive control) and the negative control. P-value (*P*) was calculated by the ‘Student’s t-test’ to determine the significance of FC of a gene per comparison. FC ≥ 2 or ≤ 0.5, and *P* < 0.05 were used as the threshold to define DEGs.

Principal component analysis (PCA) was performed with the total valid gene set (10,784 genes) as input, using R 4.1.0 software, and the ‘FactoMineR*’* and ‘Factoextra’ R packages. (29–31)

### Pathway and Gene Ontology enrichment analysis

Pathway and Gene Ontology (GO) categories analysis was performed on the Microarray Data Analysis Tool software version 3.2.0.0 (Filgen, Nagoya, Japan). Pathways were enriched using the WikiPathways system. (32)

Enrichment of both, Pathways and GO categories was performed using the gene sets (common to all three strains, or selectively shared between Colombian and Y strains) of up-regulated and down-regulated DEGs as input. P-value for each pathway and GO category was calculated with the ‘Two-tailed Fisher’s exact test’. Significantly (Z-score (*Z*) > 0, P < 0.01) enriched pathways and the top GO terms (up to ten, in order of lowest *P*) were selected for discussion. Bar plots depicting GO and pathway enrichment were generated using the ‘GOplot’ R package (33).

## Results

### Host transcriptomic remodeling per infecting *T. cruzi* strain

In the Colombian-, Y-, and Tulahuen-24 hours-infected conditions, a total of 1097 (4.9%) 1122 (5.1%), and 806 (3.6%) DEGs were identified. The proportion of up and down-regulated DEGs per condition is depicted in Figure 1. Colombian- and Y-infection led to increased numbers of up-regulated DEGs at 24-hpi. A similar trend persisted at 48-hpi but to a less evident extent. At 24-hpi, a significant overlap of up-regulated DEGs (438 of 666 (66%) in Colombian-, or of 538 (81%) in Y-infection) was observed between Colombian- and Y-infection, of which 107 (24%) were also shared with Tulahuen-infection (Figure 2, panel A). Uniquely in Tulahuen-infection, down-regulated genes over-represented the total DEGs at 24-hpi. A large proportion of down-regulated DEGs (315 of 431 (73%) in Colombian-, of 584 (54%) in Y-, or 684 (46%) in Tulahuen-infection) were shared among all infection conditions, suggesting a common pathway of cardiomyocyte functional deterioration (Figure 2, panel B).

**Figure 1.**
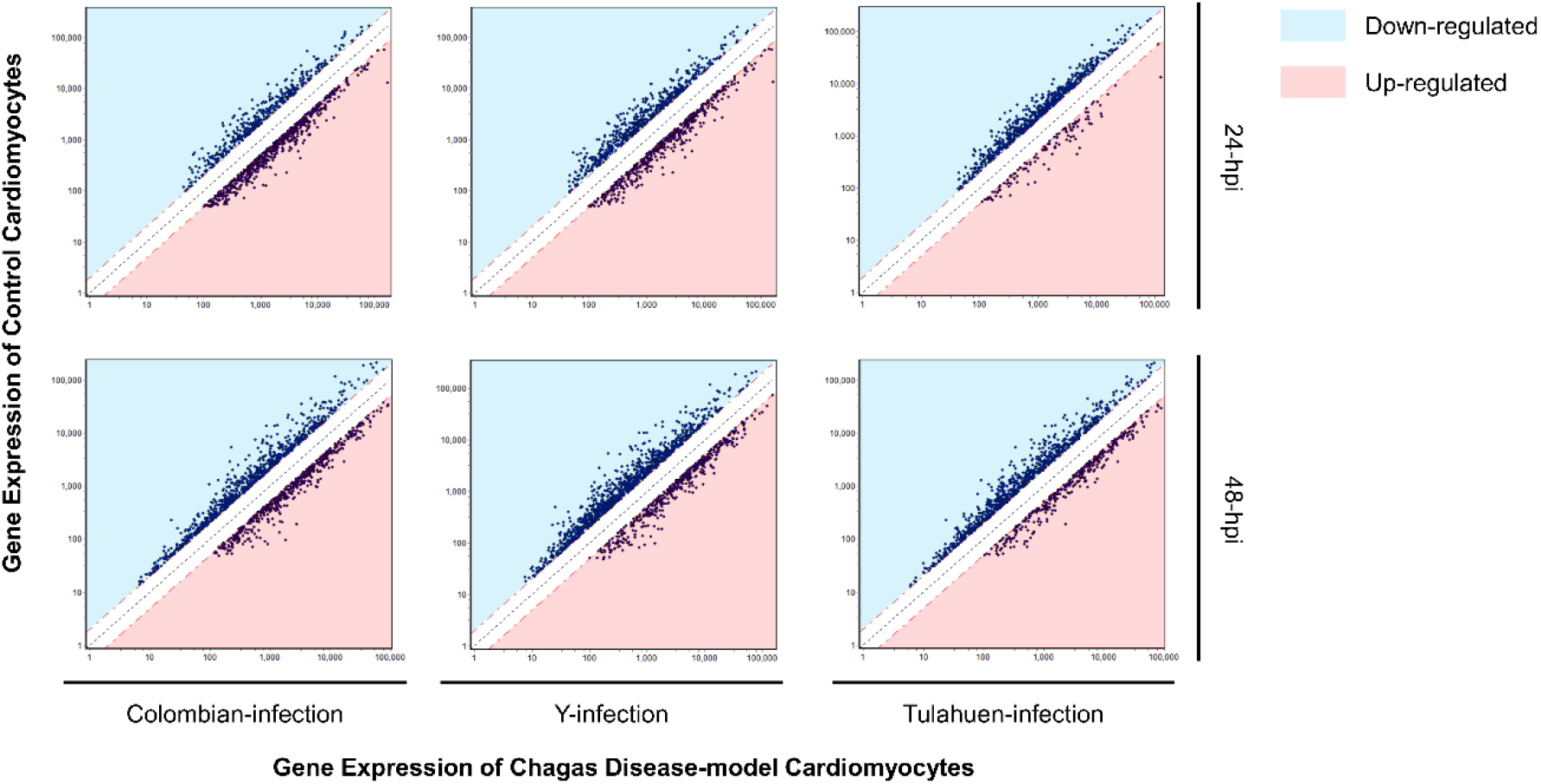
Remodeling of gene expression profiles in cardiomyocytes infected with three different *Trypanosoma cruzi* strains at two time points post-infection. Y axes indicate the gene expression level of the non-infected control cardiomyocyte, while X-axes indicate gene expression levels of *Trypanosoma cruzi*-infected Chagas cardiomyopathy model cardiomyocytes. The black dashed line, and upper and lower red dashed lines show the line of identity, and the 0.5- and 2-fold change thresholds, respectively. Each plot represents a differentially expressed gene (DEG). Blue and pink shaded areas represent down- and up-regulated DEGs, respectively. Hpi, hours post-infection.

**Figure 2.**
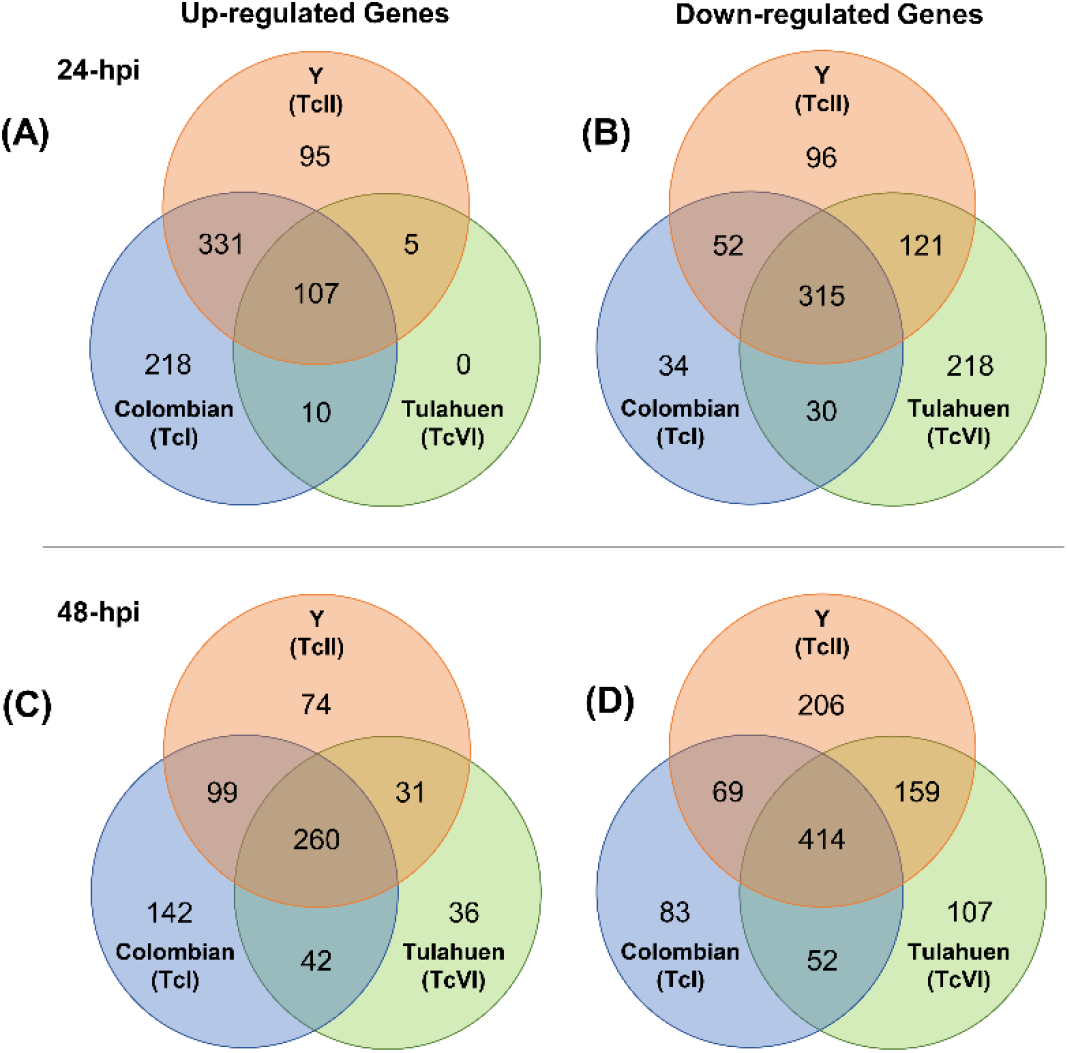
Identification of differentially expressed genes (DEGs) of the host cardiomyocyte upon *Trypanosoma cruzi* infection. Numbers of up-(left column) and down-regulated (right column) DEGs at 24- (upper row) and 48-hours post-infection (lower row) are shown in the respective Venn diagrams. Hpi, hours post-infection.

As apparent in the PCA plot, the CCM-prone Colombian- and Y-infection conditions were more closely related in host cardiomyocyte gene expression (Figure 3), potentially reflecting the significant overlap of up-regulated DEGs between the strains and the distinctive over-representation of down-regulated DEG in Tulahuen-infection. At 24-hpi, the three infection conditions showed more discrete host cardiomyocyte gene expression patterns compared with 48-hpi. This indicated that 24-hpi would serve as the more favorable time frame to investigate the uniqueness in the host transcriptional remodeling triggered by each infecting *T. cruzi* strain. Together with our interest in the earlier determinants of CCM progression in TcI/II *T. cruzi* cardiac infection, the PCA findings supported our selecting the 24-hpi stage of the *in vitro* CCM model as the main focus of the following analyses.

**Figure 3.**
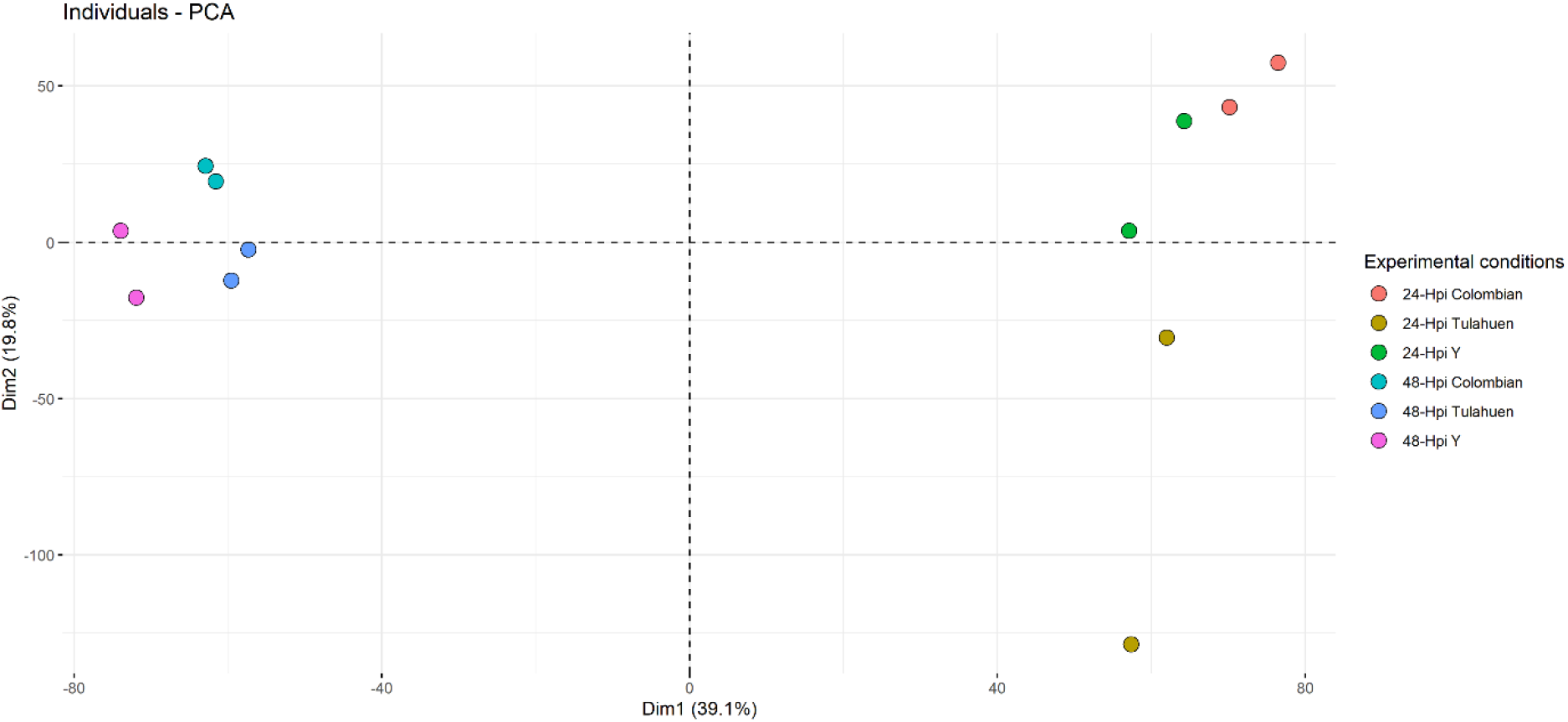
Principal component analysis plot of the transcriptional profile of the in vitro Chagas cardiomyopathy models. Closely related data points, within the infection models by different *Trypanosoma cruzi* strains at 24- and 48-hours post-infection, are visually identified by using fold change values of the total valid gene set as input. The first two principal components are plotted with their respective proportion of variance detailed within parenthesis. Hpi, hours post-infection.

### The common cardiomyocyte response to *T. cruzi* invasion

Out of the 107 commonly up-regulated genes, eight had an FC ≥ 5. The top up-regulated genes were Arresting Domain Containing4 (*Arrdc4*), Thioredoxin interacting protein (*Txnip*), and Growth/differentiation factor 15 (*Gdf15*). (Figure 4, panel A) The 107 up-regulated DEGs shared among all three infection conditions enriched pathways related to ‘Hypertrophy model’ (*Z* = 7.90, *P* < 0.001), ‘Apoptosis’ (*Z* = 4.8 *P* < 0.01), ‘Aminoacid metabolism’ (*Z* = 4.3, *P* < 0.01), ‘MAPK signaling pathway’ (*Z* = 3.9, *P* < 0.01), ‘ErbB signaling pathway’(*Z* = 4.9, *P* < 0.01), ‘Spinal cord injury’ (*Z* = 4.2, *P* < 0.01), ‘Selenium metabolism-selenoproteins’ (*Z* = 4.8, *P* < 0.01), and ‘Homologous recombination’ (*Z* = 6.5, *P* < 0.01) (Figure 5, Panel A). The same gene list was enriched in genes associated with the Biological Process GO terms of ‘Response to reactive oxygen species’ (*Z* = 8.1, *P* < 0.001), ‘Response to ionizing radiation’ (*Z* = 8.2, *P* < 0.001), ‘Response to oxidative stress’ (*Z* = 6.7, *P* < 0.001), ‘Response to hydrogen peroxide’ (*Z* = 8.1, *P* < 0.001), ‘Cellular response to stress’ (*Z* = 5.3, *P* < 0.001), ‘Primary metabolic process’ (*Z* = 3.3, *P* < 0.001), ‘Regulation of metabolic process’ (*Z* = 3.9, *P* < 0.001), and ‘Regulation of intracellular protein kinase cascade’ (*Z* = 5.4, *P* < 0.001) (Figure 5, Panel C).

**Figure 4.**
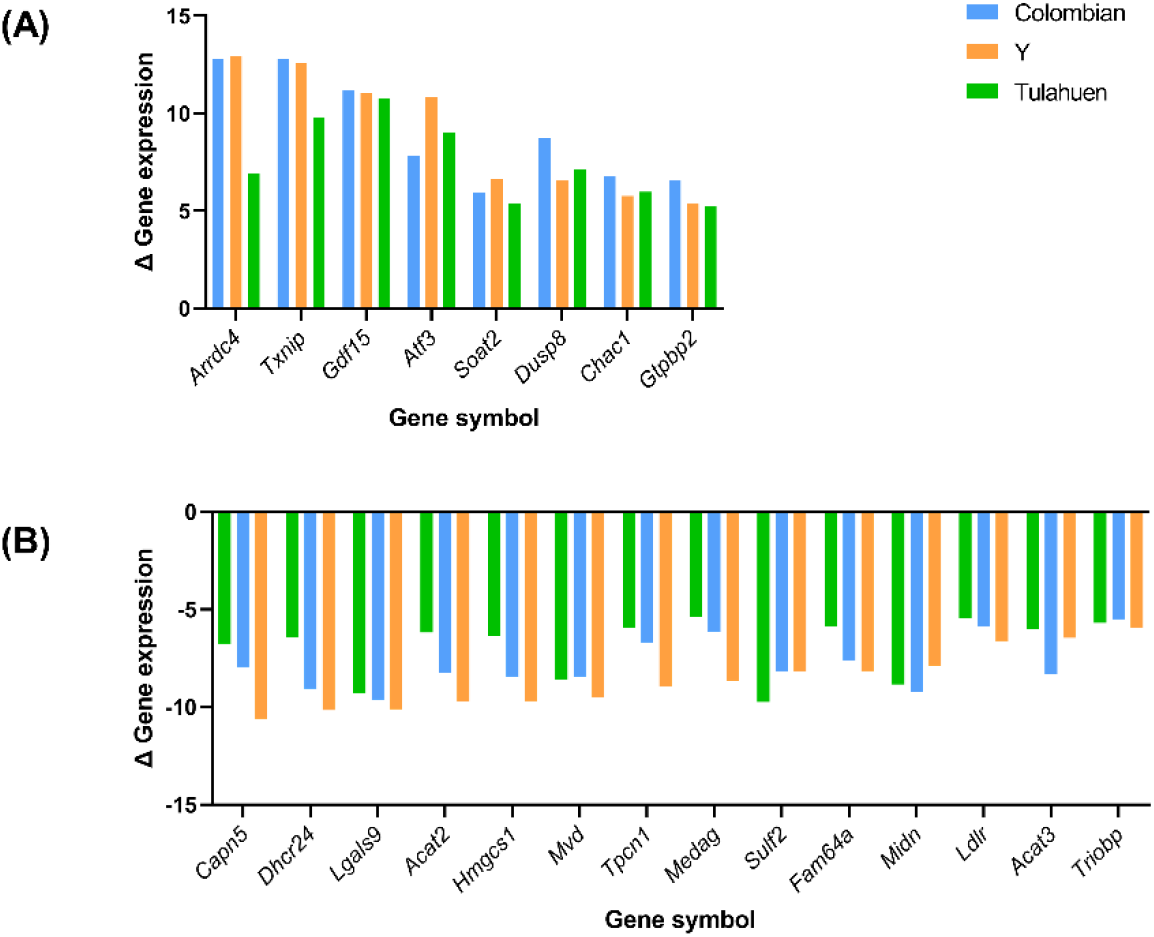
Top differentially expressed genes of the cardiomyocyte upon *Trypanosoma cruzi* infection. Extensively (A) up-regulated (fold change > 5) and (B) down-regulated (fold change < 0.2) genes of the cardiomyocyte at 24 hours post-infection are listed. Δ Gene expression, the fold change or ratio between gene expression level of the Chagas cardiomyopathy models and the non-infected negative control. *Arrdc4*, Arrestin domain containing 4; *Txnip*, Thioredoxin interacting protein; *Gdf15*, Growth differentiation factor 15; *Atf3*, Activating transcription factor 3; *Soat2*, Sterol O-acyltransferase 2; *Dusp8*, Dual specificity phosphatase 8; *Chac1*, Cation transport regulator 1; *Gtpbp2*, GTP binding protein 2; *Capn5*, Calpain 5; *Dhcr24*, 24-dehydrocholesterol reductase; *Lgals9*, Lectin, galactose binding, soluble 9; *Acat2*, Acetyl-Coenzyme A acetyltransferase 2; *Hmgcs1*, 3-hydroxy-3-methylglutaryl-Coenzyme A synthase 1; *Mvd*, Mevalonate (diphospho) decarboxylase; *Tpcn1*, Two pore channel 1; *Medag*, Mesenteric estrogen dependent adipogenesis; *Sulf2*, Sulfatase 2; *Fam64a*, Family with sequence similarity 64, member A; *Midn*, midnolin; *Ldlr*, Low density lipoprotein receptor; *Acat3*, Acetyl-Coenzyme A acetyltransferase 3; *Triobp*, TRIO and F-actin binding protein.

**Figure 5.**
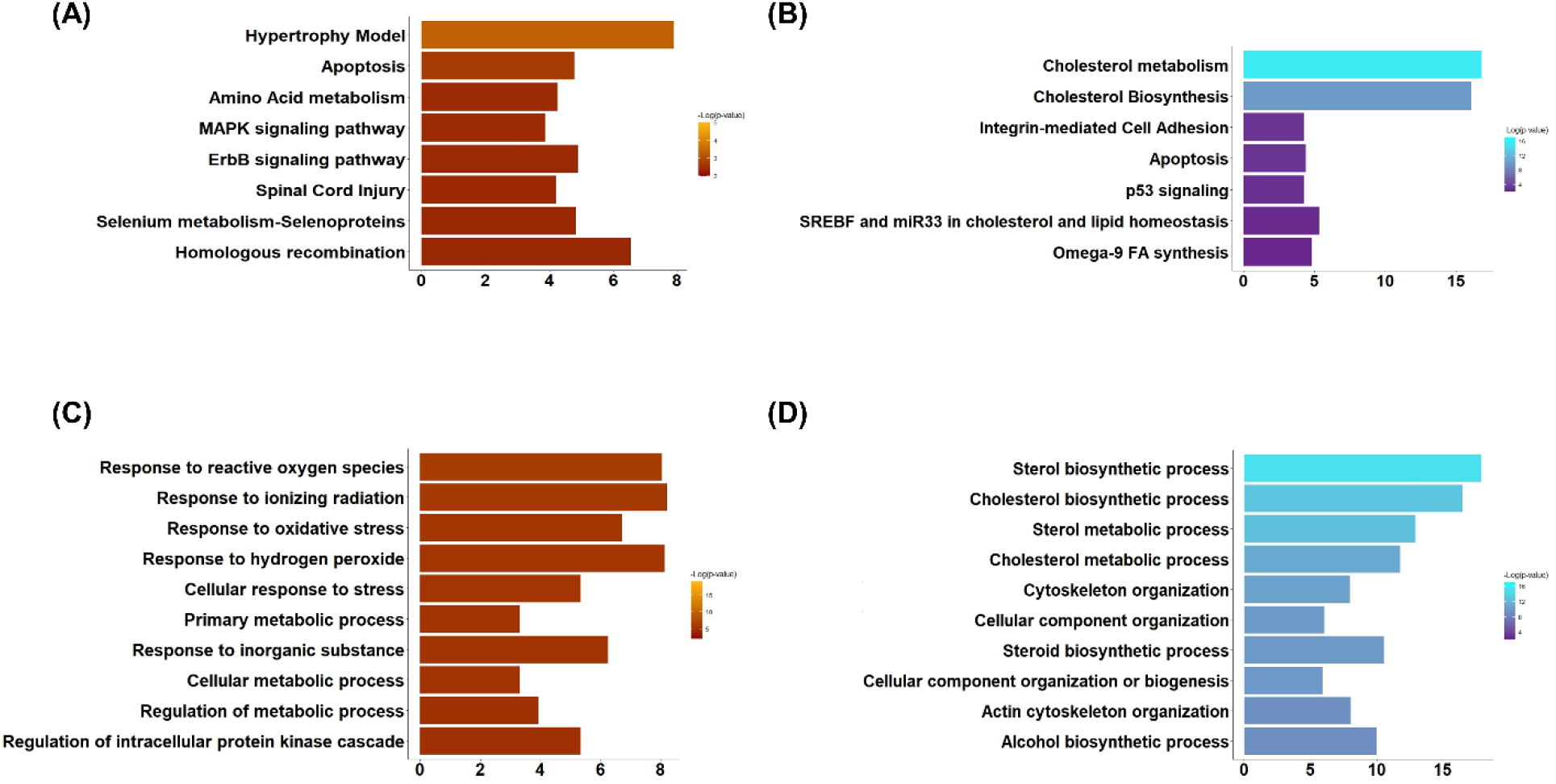
Signatures of the common cardiomyocyte response to *Trypanosoma cruzi* invasion. WikiPathways and ‘Biological Process’ Gene Ontology categories enriched by commonly (A, C) up-regulated and (B, D) down-regulated genes of the host cardiomyocyte at 24 hours post-infection are sorted based on p-value. Bar lengths represent enrichment Z-scores.

Out of the 315 commonly up-downregulated genes, fourteen had an FC ≤ 0.2 (Figure 4, Panel B). The 315 down-regulated DEGs shared among all three infection conditions enriched pathways related to ‘Cholesterol metabolism’ (*Z* = 16.8, *P* < 0.001), ‘Cholesterol biosynthesis’ (*Z* = 16.1, *P* < 0.001), ‘Integrin-mediated cell adhesion’ (*Z* = 4.2, *P* < 0.01), ‘Apoptosis’ (*Z* = 4.3, *P* < 0.01), ‘p53 signaling’ (*Z* = 4.2, *P* < 0.01), ‘SREBF and miR33 in cholesterol and lipid homeostasis’ (*Z* = 5.3, *P* < 0.01) and ‘Omega-9 FA synthesis’ (*Z* = 4.8, *P* <0.01) (Figure 5, Panel B). The same gene list was enriched in genes associated with the Biological Process GO terms of ‘Sterol biosynthetic process’ (*Z* = 17.9, *P* < 0.001), ‘Cholesterol biosynthetic process’ (*Z* = 16.5, *P* < 0.001), ‘Sterol metabolic process’ (*Z* = 12.9, *P* < 0.001), ‘Cholesterol metabolic process’ (*Z* = 11.8, *P* < 0.001), ‘Cytoskeleton organization’ (*Z* = 7.9, *P* < 0.001), ‘Cellular component organization’ (*Z* = 6.0, *P* < 0.001), ‘Steroid biosynthetic process’ (*Z* = 10.5, *P* < 0.001), ‘Cellular component organization or biogenesis’ (*Z* = 5.9, *P* < 0.001), ‘Actin cytoskeleton organization’ (*Z* = 8.1, *P* < 0.001), and ‘Alcohol biosynthetic process’ (*Z* = 10.0, *P* < 0.001) (Figure 5, Panel D).

### Transcriptomic remodeling unique to CCM-prone Colombian- and Y-infections

Up-regulated genes have over-represented the DEGs shared among the CCM-prone Colombian- and Y-infected conditions and thus formed the focus of our interest. Such 331 genes enriched pathways related to ‘DNA replication (*Z* = 6.8, *P* < 0.001)’, ‘One Carbon metabolism (*Z* = 6.8, *P* < 0.001)’, ‘Nucleotide metabolism (*Z* = 6.8, *P* < 0.001)’, ‘Glutathione and one-carbon metabolism (*Z* = 5.1, *P* < 0.01)’, and ‘Mitochondrial gene expression (*Z* = 4.9, *P* < 0.01)’ (Figure 6, Panel A). The same gene list was enriched in genes associated with the Biological Process GO terms of ‘Nucleic acid metabolic process’ (*Z* = 8.3, *P* < 0.001), ‘Nucleobase-containing compound metabolic process (*Z* = 7.2, *P* < 0.001), ‘Heterocycle metabolic process’ (*Z* = 7.1, *P* < 0.001), ‘Cellular nitrogen compound metabolic process’ (*Z* = 6.9, *P* < 0.001), ‘Cellular aromatic compound metabolic process’ (*Z* = 6.9, *P* < 0.001), ‘Organic cyclic compound metabolic process’ (*Z* = 6.8, *P* < 0.001), ‘Nitrogen compound metabolic process’ (*Z* = 6.7, *P* < 0.001), ‘RNA metabolic process’ (*Z* = 6.9, *P* < 0.001), and ‘Gene expression’ (*Z* = 6.1, *P* < 0.001) (Figure 6, Panel B).

**Figure 6.**
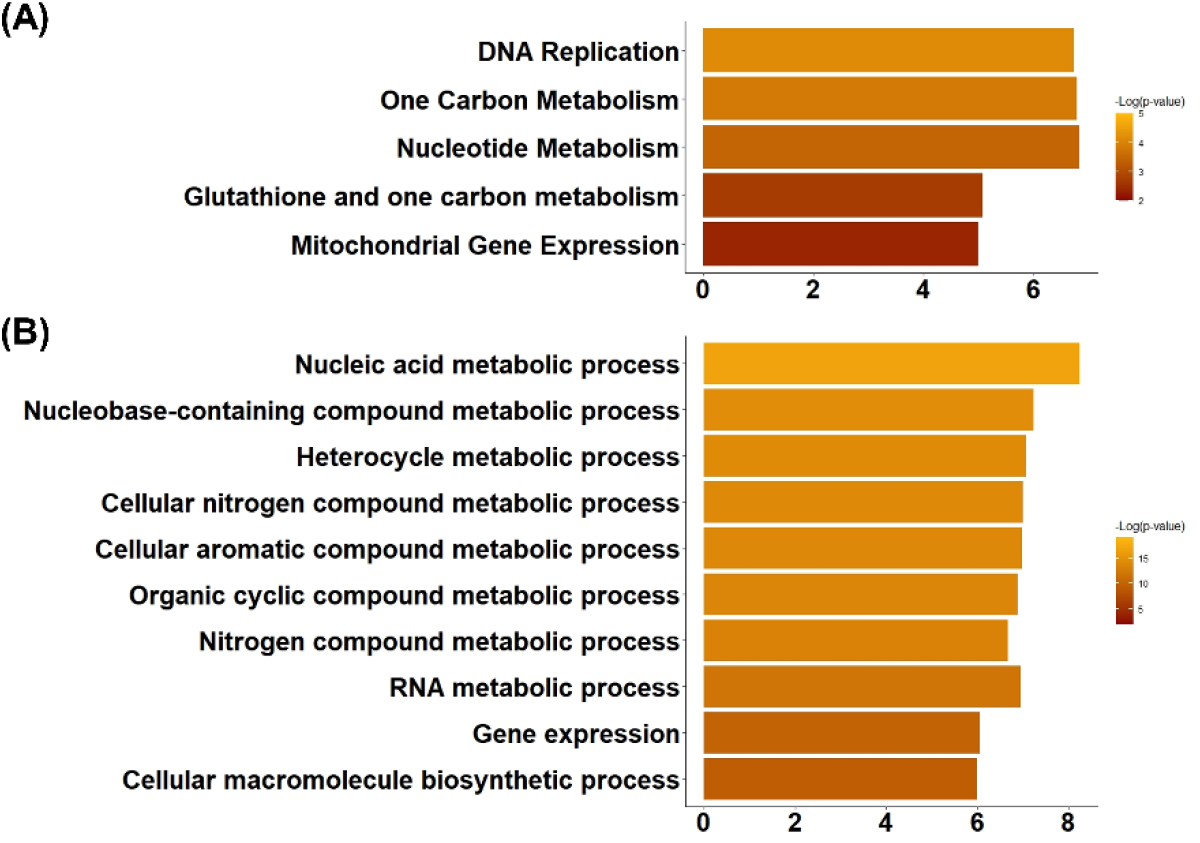
Signatures of transcriptomic remodeling unique to cardiomyopathy-prone Colombian- and Y-infections. WikiPathways enriched by the genes uniquely up- regulated at 24 hours post-infection in Colombian- and Y-infected Chagas cardiomyopathy models are sorted based on p-value. Bar lengths represent enrichment Z-scores.

Differentially expressed genes corresponding to the ‘Glutathione and one-carbon metabolism’ pathway (Wikipathway ID: WP730) were individually tracked and analyzed, where the genes corresponding to the ‘one-carbon metabolism’ and ‘Glutathione metabolism’ showed an up and downregulation trend, respectively. Differences on the Pathways and GO terms that are enriched with the genes that are upregulated in the cardiomyocytes as a response against infection with each one of the *T. cruzi* strains versus the CCM-prone TcI/II are summarized in Table 1.

**Table 1.**
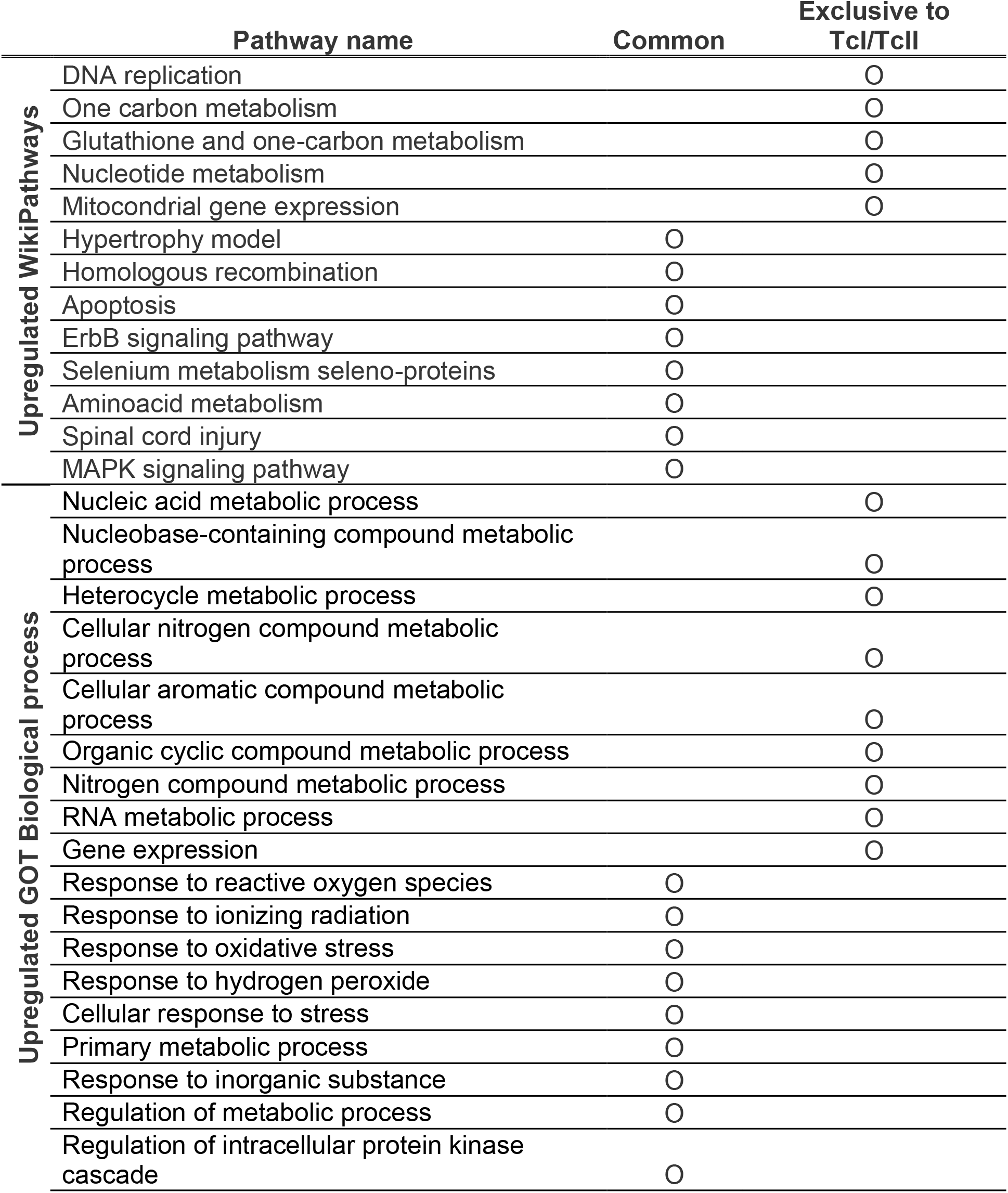
Summarization of the pathways enriched by the upregulated genes in all the *T. Cruzi-infection* conditions.

### Transcriptomic remodeling unique to TcVI Tulahuen-infections

The digestive-form-causing TcVI Tulahuen-infection led to unique down-regulation of 218 DEGs in the cardiomyocyte transcriptome. Significantly enriched pathways were: ‘Proteosome degradation’ (*Z* = 5.1, *P* < 0.001), ‘Electron transport chain’ (*Z* = 4.1, *P* < 0.01), and ‘Oxidative phosphorylation’ (*Z* = 3.7, *P* < 0.01). Top 10 Biological processes GO terms enriched by this gene set are ‘Cellular metabolic process’ (*Z* = 3.9, *P* < 0.001), ‘Metabolic process’ (*Z* = 3.4, *P* < 0.001), ‘Cellular process’ (*Z* = 2.3, *P* < 0.001), ‘Organic substance metabolic process’ (*Z* = 3.2, *P* < 0.001), ‘Localization’ (*Z* = 4.0, *P* < 0.001), ‘Organic substance transport’ (*Z* = 4.5, *P* < 0.001), ‘Macromolecule localization’ (*Z* = 4.4, *P* < 0.001), ‘Cellular protein metabolic process’ (*Z* = 3.9, *P* < 0.001), and ‘Cellular protein metabolic process’ (*Z* = 3.9, *P* < 0.001).

## Discussion

*In vitro* CCM modeling, using HL-1 cardiomyocytes and *T. cruzi* from different genetic strains, and the subsequent transcriptomic analyses, have characterized in depth the myocardial innate defense response that are potential determinants of CCM-progression.

Firstly, the cross-strain comparative analysis has led to the characterization of the universal hallmarks of the cardiomyocyte defense response against *T. cruzi* infection, consisting of DEGs evoked upon all three infection conditions. Among the pathways and GO terms enriched by such hallmark, transcriptomic signatures were those related to cardiac hypertrophy (the ‘Hypertrophy model’ and ‘MAPK signaling’ pathways) and heart energy metabolism (the ‘SREBF and miR33 in cholesterol and lipid homeostasis’, and ‘Omega-9 FA synthesis’ pathways and the ‘Sterol biosynthetic process’, ‘Cholesterol biosynthetic process’, ‘Sterol metabolic process’, ‘Cholesterol metabolic process’, ‘Steroid biosynthetic process’ GO terms). Upregulation of ErbB signaling, as well as its downstream MAPK pathway (34), are promotive of the cardiomyocyte hypertrophic response. While initially adaptive to stress stimuli, such signaling pathways are the underlying cause of pathological cardiac hypertrophy and mechanical dysfunction when chronically activated. Thus, these pathways, centralizing in the adaptive/maladaptive myocardial hypertrophic response, have constituted the ‘double-edged’ roles in the development of, and the common potential targets for, ischemic and viral cardiomyopathies (37–39). Cellular metabolism can drastically influence cellular function. A switching from beta-oxidation to glycolysis as the major source of energy production, resembling the Warburg effect, has been known as the cardinal metabolic feature of the failing heart (35,36). Here, *Arrdc4*, having been most extensively modulated at the transcriptional level in our *in vitro* CCM models, may cause glucose deprivation in the cardiomyocytes leading to cell death (37). Likewise, *Txnip* will inhibit cellular glucose uptake through its inhibitory action towards thioredoxin (38,39). Together with such transcriptional profiles altogether leading towards dampened glycolysis, we also observed evidence of lipid metabolism imbalance. The uncoupling of lipid synthesis and beta-oxidation has been reported as a hallmark metabolic feature in CCM (51). Sterol regulatory element binding factor (*Srebf*), a transcription factor that regulates cellular lipid homeostasis by controlling the expression of an array of enzymes required for endogenous cholesterol, fatty acid, triacylglycerol, and phospholipid synthesis (40,41), has been proposed as the potential target for ameliorating the cardiac lipidopathy in Chagas disease (54, 55). This unbalance of lipid synthesis and beta-oxidation became further prominent in our *in vitro* CCM model at 48-hpi, accompanied by the down regulation of ‘Fatty acid oxidation’ and ‘Beta-oxidation’ pathway-related genes: *Acss2, Acsl4*, and *Acsl5* (data not shown) (54). The correction of such pathogenic metabolic uncoupling that may even potentiate *T. cruzi* epimastigotes proliferation, together with counteracting the energetically unfavorable cardiac Warburg effect, may exert therapeutic potentials against CCM through the restoration of metabolic homeostasis (42–44).

Secondly, and most interestingly, the CCM-prone Colombian- and Y-infections led to the more aggressive transcriptomic remodeling of the cardiomyocyte at 24-hpi, compared with the TcVI Tulahuen-infection. Furthermore, the shared signatures of the cardiomyocyte response against these two strains highlighted an intriguing role of glutathione metabolism and Reactive Oxygen Species (ROS) mitigation in CCM-progression. ROS mitigation is, not a unique host response to *T. cruzi* cardiac infection, but in fact a shared defense mechanism of the cardiomyocyte against the more common insults leading to dilated cardiomyopathy, namely ischemia/hypoxia and coxsackie virus-infection (45–47). Previous *in vitro* (26,27) and *in vivo* studies (28) have shown that glutathione and its related metabolites serve as key antioxidants of the cardiomyocyte under infectious or hypoxic stress (43–46). In the present study, the derangement in cardiomyocyte ROS homeostasis, represented by the gene enrichment in the ‘Selenium metabolism-selenoproteins’ pathway and the ‘Response to reactive oxygen species’, ‘Response to ionizing radiation’, ‘Response to oxidative stress’, and ‘Response to hydrogen peroxide’ GO terms upon *T. cruzi* infection, was further accentuated by the CCM-prone strains, leading to significant upregulation of the compensatory ‘Glutathione and one-carbon metabolism’ pathways in infected cardiomyocytes.

Additionally, among the top up-regulated DEGs was the TGF-beta superfamily *Gdf15*, an emerging cardiovascular biomarker. GDF15 is known to protect cardiomyocytes from ROS and nitric oxide-induced apoptosis *in vitro*. Also, the risk stratification potential of GDF15 in heart failure patients is currently gaining attention (51–53). Now appreciating that the upregulation in GDF15 comprises the cardinal feature of cardiomyocyte response in our *in vitro* CCM model, the behavior of this cytokine in actual CCM patients arises as our next intriguing research question.

In conclusion, on top of the ubiquitously observed hypertrophic and metabolic remodeling in response to *T. cruzi* infection, the cardio-virulent strains elicited an accentuated defect in cardiomyocyte ROS mitigation, suggesting its underpinning role in CCM progression.

## References

1. Chagas C. Nova tripanozomiaze humana: estudos sobre a morfolojia e o ciclo evolutivo do Schizotrypanum cruzi n. gen., n. sp., ajente etiolojico de nova entidade morbida do homem. Mem Inst Oswaldo Cruz [Internet]. 1909 Aug [cited 2022 Dec 26];1(2):159–218. Available from: http://www.scielo.br/j/mioc/a/FXTzX4Sptjs5Wnz3yKCMRpS/abstract/?lang=pt

2. Chagas disease - PAHO/WHO | Pan American Health Organization [Internet]. [cited 2022 Dec 26]. Available from: https://www.paho.org/en/topics/chagas-disease

3. Lidani KCF, Andrade FA, Bavia L, Damasceno FS, Beltrame MH, Messias-Reason IJ, et al. Chagas disease: From discovery to a worldwide health problem. J Phys Oceanogr. 2019 Jul 2;49(6):166.

4. Nunes MCP, Beaton A, Acquatella H, Bern C, Bolger AF, Echeverría LE, et al. Chagas Cardiomyopathy: An Update of Current Clinical Knowledge and Management: A Scientific Statement From the American Heart Association. Circulation [Internet]. 2018 Sep 18 [cited 2022 Dec 26];138(12):e169–209. Available from: https://www.ahajournals.org/doi/abs/10.1161/CIR.0000000000000599

5. Stride N, Larsen S, Hey-Mogensen M, Hansen CN, Prats C, Steinbrüchel D, et al. Impaired mitochondrial function in chronically ischemic human heart. Am J Physiol Heart Circ Physiol [Internet]. 2013 [cited 2022 Dec 26];304(11):1407–14. Available from: http://www.ajpheart.org

6. Knowlton KU. Dilated Cardiomyopathy: Viral Persistence and a Role for a Viral Proteinase in Human Hearts. Circulation [Internet]. 2019 May 14 [cited 2022 Dec 26];139(20):2339–41. Available from: https://www.ahajournals.org/doi/abs/10.1161/CIRCULATIONAHA.119.040037

7. Bouin A, Gretteau PA, Wehbe M, Renois F, N’Guyen Y, Lévêque N, et al. Enterovirus Persistence in Cardiac Cells of Patients with Idiopathic Dilated Cardiomyopathy Is Linked to 5’ Terminal Genomic RNA-Deleted Viral Populations with Viral-Encoded Proteinase Activities. Circulation [Internet]. 2019 May 14 [cited 2022 Dec 26];139(20):2326–38. Available from: https://www.ahajournals.org/doi/abs/10.1161/CIRCULATIONAHA.118.035966

8. Bayeva M, Teodor K. Molecular and Cellular Basis of Viable Dysfunctional Myocardium. 2014 [cited 2022 Dec 26]; Available from: http://circheartfailure.ahajournals.org

9. Page BJ, Young RF, Suzuki G, Fallavollita JA, Canty JM. The physiological significance of a coronary stenosis differentially affects contractility and mitochondrial function in viable chronically dysfunctional myocardium. Basic Res Cardiol. 2013;108(4).

10. Hershberger RE, Pinto JR, Parks SB, Kushner JD, Li D, Ludwigsen S, et al. Clinical and Functional Characterization of TNNT2 Mutations Identified in Patients With Dilated Cardiomyopathy. Circ Cardiovasc Genet [Internet]. 2009 Aug [cited 2023 Jan 8];2(4):306–13. Available from: https://www.ahajournals.org/doi/abs/10.1161/CIRCGENETICS.108.846733

11. Poller W, Fechner H, Noutsias M, Tschoepe C, Pauschinger M, Schultheiss HP. The molecular basis of cardiotropic viral infections. European Heart Journal Supplements [Internet]. 2002 Dec 1 [cited 2023 Jan 10];4(Suppl_I):I18–30. Available from: https://academic.oup.com/eurheartjsupp/article/4/suppl_I/I18/442308

12. Rao L, Debbas M, Sabbatini P, Hockenbery D, Korsmeyer S, White E. The adenovirus E1A proteins induce apoptosis, which is inhibited by the E1B 19-kDa and Bcl-2 proteins. Proc Natl Acad Sci U S A [Internet]. 1992 [cited 2023 Jan 23];89(16):7742–6. Available from: https://pubmed.ncbi.nlm.nih.gov/1457005/

13. Kawai C. From Myocarditis to Cardiomyopathy: Mechanisms of Inflammation and Cell Death. Circulation [Internet]. 1999 Mar 2 [cited 2023 Jan 23];99(8):1091–100. Available from: https://www.ahajournals.org/doi/abs/10.1161/01.cir.99.8.1091

14. Gougeon ML, Montagnier L. Apoptosis in AIDS. Science. 1993;260.

15. Debiasi RL, Robinson BA, Leser JS, Brown RD, Long CS, Clarke P. Critical role for Death-Receptor Mediated Apoptotic Signaling in Viral Myocarditis. J Card Fail [Internet]. 2010 Nov [cited 2023 Jan 23];16(11):901. Available from: /pmc/articles/PMC2994069/

16. Zingales B, Miles MA, Campbell DA, Tibayrenc M, Macedo AM, Teixeira MMG, et al. The revised Trypanosoma cruzi subspecific nomenclature: rationale, epidemiological relevance and research applications. Infect Genet Evol [Internet]. 2012 Mar [cited 2022 Dec 26];12(2):240–53. Available from: https://pubmed.ncbi.nlm.nih.gov/22226704/

17. Poveda C, Fresno M, Ria Gironè S N, Martins-Filho OA, David Ramírez J, Santi-Rocca J, et al. Cytokine Profiling in Chagas Disease: Towards Understanding the Association with Infecting Trypanosoma cruzi Discrete Typing Units (A BENEFIT TRIAL Sub-Study). [cited 2022 Dec 26]; Available from: http://www.plosone.org

18. Ramírez JD, Guhl F, Rendón LM, Rosas F, Marin-Neto JA, Morillo CA. Chagas cardiomyopathy manifestations and Trypanosoma cruzi genotypes circulating in chronic Chagasic patients. PLoS Negl Trop Dis [Internet]. 2010 Nov [cited 2022 Dec 26];4(11). Available from: https://pubmed.ncbi.nlm.nih.gov/21152056/

19. Zafra G, Mantilla JC, Valadares HM, Macedo AM, González CI. Evidence of Trypanosoma cruzi II infection in Colombian chagasic patients. Parasitol Res [Internet]. 2008 Aug [cited 2022 Dec 26];103(3):731–4. Available from: https://pubmed.ncbi.nlm.nih.gov/18523803/

20. Burgos JM, Diez M, Vigliano C, Bisio M, Risso M, Duffy T, et al. Molecular identification of Trypanosoma cruzi discrete typing units in end-stage chronic Chagas heart disease and reactivation after heart transplantation. Clin Infect Dis [Internet]. 2010 Sep 1 [cited 2022 Dec 26];51(5):485–95. Available from: https://pubmed.ncbi.nlm.nih.gov/20645859/

21. Añez N, Crisante G, da Silva FM, Rojas A, Carrasco H, Umezawa ES, et al. Predominance of lineage I among Trypanosoma cruzi isolates from Venezuelan patients with different clinical profiles of acute Chagas’ disease. Trop Med Int Health [Internet]. 2004 [cited 2022 Dec 26];9(12):1319–26. Available from: https://pubmed.ncbi.nlm.nih.gov/15598264/

22. Nielebock MAP, Moreira OC, das Chagas Xavier SC, de Freitas Campos Miranda L, de Lima ACB, de Jesus Sales Pereira TO, et al. Association between Trypanosoma cruzi DTU TcII and chronic Chagas disease clinical presentation and outcome in an urban cohort in Brazil. PLoS One [Internet]. 2020 Dec 1 [cited 2022 Dec 26];15(12). Available from: /pmc/articles/PMC7710061/

23. Cura CI, Lucero RH, Bisio M, Oshiro E, Formichelli LB, Burgos JM, et al. Trypanosoma cruzi Discrete Typing Units in Chagas disease patients from endemic and non-endemic regions of Argentina. Parasitology [Internet]. 2012 Apr [cited 2022 Dec 26];139(4):516–21. Available from: https://www.cambridge.org/core/journals/parasitology/article/abs/trypanosoma-cruzi-discrete-typing-units-in-chagas-disease-patients-from-endemic-and-nonendemic-regions-of-argentina/8AFB9E04537CEB3E797A706FD6690CE5

24. Monje-Rumi MM, Floridia-Yapur N, Zago MP, Ragone PG, Pérez Brandán CM, Nuñez S, et al. Potential association of Trypanosoma cruzi DTUs TcV and TcVI with the digestive form of Chagas disease. Infection, Genetics and Evolution. 2020 Oct 1;84:104329.

25. Castro TBR de, Canesso MCC, Boroni M, Chame DF, Souza D de L, Toledo NE de, Birelli Tahara E, Pena SD, Machado SR, Chiari E, Macedo A, Franco GR. Differential Modulation of Mouse Heart Gene Expression by Infection With Two Trypanosoma cruzi Strains: A Transcriptome Analysis. Front Genet. 2020 Sep 3;11:1031.

26. Adesse D, Garzoni LR, Meirelles MDN, Iacobas DA, Iacobas S, Spray DC. Transcriptomic signatures of alterations in a myoblast cell line infected with four distinct strains of Trypanosoma cruzi. Am J Trop Med Hyg [Internet]. 2010 May [cited 2022 Dec 2];82(5):846–54. Available from: https://pubmed.ncbi.nlm.nih.gov/20439965/

27. Nisimura LM, Coelho LL, de Melo TG, Vieira P de C, Victorino PH, Garzoni LR, Spray DC, Iacobas DA, Iacobas S, Tanowitz HB, Adesse D. Trypanosoma cruzi Promotes Transcriptomic Remodeling of the JAK/STAT Signaling and Cell Cycle Pathways in Myoblasts. Front Cell Infect Microbiol. 2020 Jun 17;10:255.

28. Contreras VT, Salles JM, Thomas N, Morel CM, Goldenberg S. In vitro differentiation of Trypanosoma cruzi under chemically defined conditions. Mol Biochem Parasitol. 1985 Sep 1;16(3):315–27.

29. R core Team. R: A Language and Environmentfor Statistical Computing. [Internet]. Vienna, Asutria; 2022. Available from: https://www.r-project.org/.

30. Lê S, Josse J, Husson F. FactoMineR: An R Package for Multivariate Analysis. J Stat Softw [Internet]. 2008 Mar 18 [cited 2023 Jan 27];25(1):1–18. Available from: https://www.jstatsoft.org/index.php/jss/article/view/v025i01

31. Kassambara A, Mundt F. factoextra: Extract and Visualize the Results of Multivariate Data Analyses. [Internet]. 2020. Available from: https://cran.r-project.org/package=factoextra

32. Martens M, Ammar A, Riutta A, Waagmeester A, Slenter DN, Hanspers K, et al. WikiPathways: connecting communities. Nucleic Acids Res [Internet]. 2021 Jan 8 [cited 2023 Jan 27];49(D1):D613–21. Available from: https://pubmed.ncbi.nlm.nih.gov/33211851/

33. Walter W, Sánchez-Cabo F, Ricote M. GOplot: an R package for visually combining expression data with functional analysis. Bioinformatics [Internet]. 2015 Feb 6 [cited 2023 Jan 27];31(17):2912–4. Available from: https://pubmed.ncbi.nlm.nih.gov/25964631/

34. Rohrbach S, Yan X, Weinberg EO, Hasan F, Bartunek J, Marchionni MA, et al. Neuregulin in Cardiac Hypertrophy in Rats With Aortic Stenosis. Circulation [Internet]. 1999 Jul 27 [cited 2023 Jan 4];100(4):407–12. Available from: https://www.ahajournals.org/doi/abs/10.1161/01.CIR.100.4.407

35. Taylor LA, Carthy CM, Yang D, Saad K, Wong D, Schreiner G, et al. Host Gene Regulation During Coxsackievirus B3 Infection in Mice. Circ Res [Internet]. 2000 Aug 18 [cited 2023 Jan 13];87(4):328–34. Available from: https://www.ahajournals.org/doi/abs/10.1161/01.res.87.4.328

36. Abe H, Takeda N, Isagawa T, Semba H, Nishimura S, Suimye Morioka M, et al. Macrophage hypoxia signaling regulates cardiac fibrosis via Oncostatin M. [cited 2023 Feb 17]; Available from: https://doi.org/10.1038/s41467-019-10859-w

37. Nakayama Y, Mukai N, Kreitzer G, Patwari P, Yoshioka J. Interaction of ARRDC4 With GLUT1 Mediates Metabolic Stress in the Ischemic Heart. Circ Res [Internet]. 2022 Sep 2 [cited 2022 Dec 14];131(6):510–27. Available from: https://www.ahajournals.org/doi/abs/10.1161/CIRCRESAHA.122.321351

38. Patwari P, Chutkow WA, Cummings K, Verstraeten VLRM, Lammerding J, Schreiter ER, et al. Thioredoxin-independent regulation of metabolism by the alpha-arrestin proteins. J Biol Chem [Internet]. 2009 Sep 11 [cited 2022 Dec 13];284(37):24996–5003. Available from: https://pubmed.ncbi.nlm.nih.gov/19605364/

39. Kaimul AM, Nakamura H, Masutani H, Yodoi J. Thioredoxin and thioredoxin-binding protein-2 in cancer and metabolic syndrome. Free Radic Biol Med. 2007 Sep 15;43(6):861–8.

40. Horton JD, Goldstein JL, Brown MS. SREBPs: activators of the complete program of cholesterol and fatty acid synthesis in the liver. J Clin Invest [Internet]. 2002 May 1 [cited 2022 Dec 13];109(9):1125–31. Available from: https://pubmed.ncbi.nlm.nih.gov/11994399/

41. Eberlé D, Hegarty B, Bossard P, Ferré P, Foufelle F. SREBP transcription factors: master regulators of lipid homeostasis. Biochimie [Internet]. 2004 [cited 2022 Dec 13];86(11):839–48. Available from: https://pubmed.ncbi.nlm.nih.gov/15589694/

42. Pereira MG, Nakayasu ES, Sant’Anna C, de Cicco NNT, Atella GC, de Souza W, et al. Trypanosoma cruzi Epimastigotes Are Able to Store and Mobilize High Amounts of Cholesterol in Reservosome Lipid Inclusions. PLoS One [Internet]. 2011 [cited 2022 Dec 13];6(7):e22359. Available from: https://journals.plos.org/plosone/article?id=10.1371/journal.pone.0022359

43. Pereira MG, Visbal G, Costa TFR, Frases S, de Souza W, Atella G, et al. Trypanosoma cruzi epimastigotes store cholesteryl esters in lipid droplets after cholesterol endocytosis. Mol Biochem Parasitol. 2018 Sep 1;224:6–16.

44. Johndrow C, Nelson R, Tanowitz H, Weiss LM, Nagajyothi F. Trypanosoma cruzi infection results in an increase in intracellular cholesterol. Microbes Infect. 2014 Apr 1;16(4):337–44.

45. Machado FS, Tanowitz HB, Ribeiro AL. Pathogenesis of Chagas cardiomyopathy: Role of inflammation and oxidative stress. J Am Heart Assoc [Internet]. 2013 [cited 2022 Dec 13];2(5). Available from: http://ahajournals.org

46. Tschöpe C, Ammirati E, Bozkurt B, Caforio ALP, Cooper LT, Felix SB, et al. Myocarditis and inflammatory cardiomyopathy: current evidence and future directions. Nature Reviews Cardiology 2020 18:3 [Internet]. 2020 Oct 12 [cited 2022 Dec 16];18(3):169–93. Available from: https://www.nature.com/articles/s41569-020-00435-x

47. Tsutsui H, Kinugawa S, Matsushima S. Oxidative stress and heart failure. Am J Physiol Heart Circ Physiol [Internet]. 2011 Dec [cited 2022 Dec 16];301(6):2181–90. Available from: https://journals.physiology.org/doi/10.1152/ajpheart.00554.2011

48. Pérez-Fuentes R, Guégan JF, Barnabé C, López-Colombo A, Salgado-Rosas H, Torres-Rasgado E, et al. Severity of chronic Chagas disease is associated with cytokine/antioxidant imbalance in chronically infected individuals. Int J Parasitol. 2003 Mar 1;33(3):293–9.

49. Wen J jun, Yachelini PC, Sembaj A, Manzur RE, Garg NJ. Increased oxidative stress is correlated with mitochondrial dysfunction in chagasic patients. Free Radic Biol Med. 2006 Jul 15;41(2):270–6.

50. de Oliveira TB, Pedrosa RC, Filho DW. Oxidative stress in chronic cardiopathy associated with Chagas disease. Int J Cardiol. 2007 Apr 4;116(3):357–63.

51. Wesseling M, de Poel JHC, de Jager Sca. Growth differentiation factor 15 in adverse cardiac remodelling: from biomarker to causal player. ESC Heart Fail [Internet]. 2020 Aug 1 [cited 2023 Jan 7];7(4):1488. Available from: /pmc/articles/PMC7373942/

52. Heger J, Schiegnitz E, von Waldthausen D, Anwar MM, Piper HM, Euler G. Growth differentiation factor 15 acts anti-apoptotic and pro-hypertrophic in adult cardiomyocytes. J Cell Physiol [Internet]. 2010 Jul [cited 2023 Jan 7];224(1):120–6. Available from: https://pubmed.ncbi.nlm.nih.gov/20232299/

53. Xu J, Kimball TR, Lorenz JN, Brown DA, Bauskin AR, Klevitsky R, et al. GDF15/MIC-1 Functions As a Protective and Antihypertrophic Factor Released From the Myocardium in Association With SMAD Protein Activation. Circ Res [Internet]. 2006 Feb 17 [cited 2022 Dec 14];98(3):342–50. Available from: https://www.ahajournals.org/doi/abs/10.1161/01.RES.0000202804.84885.d0

